# ERGA-BGE genome of the Spanish Moon Moth *Graellsia isabellae* Graells, 1849: a nocturnal lepidopteran protected by the Habitats Directive

**DOI:** 10.1101/2025.01.08.631683

**Authors:** Marta Vila, Neus Marí-Mena, Roger Vila, Ana Riesgo, Daniel García-Souto, Carlos Lopez-Vaamonde, Astrid Böhne, Rita Monteiro, Thomas Marcussen, Rebekah Oomen, Torsten Hugo Struck, Marta Gut, Laura Aguilera, Francisco Câmara Ferreira, Fernando Cruz Rodriguez, Jèssica Gómez-Garrido, Tyler S. Alioto, Chiara Bortoluzzi

## Abstract

The reference genome of the Spanish Moon Moth, *Graellsia isabellae*, will be of great importance for evolutionary and conservation genomics. Firstly, this reference genome, alongside phylogenomic analyses, may finally resolve the longstanding debate regarding the scientific name of this iconic species—whether it should be *Graellsia isabellae* (Graells, 1849) or *Actias isabellae* (Graells, 1849). Secondly, the reference genome will be instrumental in the genetic monitoring of this protected species, enabling advanced methods to calculate contemporary population genomics estimates. The genome was assembled into 31 contiguous chromosomal pseudomolecules (Z chromosome included). The mitochondrial genome has also been assembled and is 15,247 bp in length. This chromosome-level assembly encompasses 0.56 Gb, composed of 38 contigs and 32 scaffolds, with contig and scaffold N50 values of 18.9 Mb and 20.4 Mb, respectively.

## Introduction

The Spanish Moon Moth, *Graellsia isabellae* (Graells, 1849), also referred to as *Actias isabellae* (Graells, 1849), is a species of lepidopteran in the family *Saturniidae* with a diploid number of 2n = 62 chromosomes according to karyotype analysis (Templado et al. 1973). This univoltine insect is primarily found in the mountainous regions of the Eastern Iberian Peninsula and the French Alps. The larvae predominantly feed on Scots pine (*Pinus sylvestris* L.), although some populations in Spain utilise Black pine (*P. nigra* Arnold). Marí-Mena et al. (2016) identified a strong phylogeographic pattern, revealing six genetic groups distributed mainly along distinct mountain ranges, though no host-associated differentiation was detected. The nuclear microsatellite markers used in that work (Vila et al. 2010) also revealed a west–east cline in allele frequencies along the Central Pyrenees that causes low overall genetic differentiation within and around the Ordesa y Monte Perdido National Park (González-Castellano et al. 2023).

The Spanish Moon Moth is protected under the Bern Convention (ETS No. 104, Annex III), as well as the European Union’s Habitats Directive (92/43/EEC, Annexes II and V). Consequently, Spain and France, where the species occurs naturally, have implemented conservation measures aimed at preventing any disturbance to the moth or its habitats. *Graellsia isabellae* is currently classified as “Data Deficient” on the IUCN Red List. This classification was assigned in 1996 and is due for an update. Marí-Mena et al. (2019) suggested reclassifying it as “Least Concern”, reflecting a generally favorable conservation status. However, Marí-Mena et al. (2019) also cautioned that some findings, such as a high temporal fragmentation index and very low values of *N*_e_*/N*, suggest a potential risk of genetic erosion if populations become isolated due to habitat fragmentation.

The generation of this reference resource was coordinated by the European Reference Genome Atlas (ERGA) initiative’s Biodiversity Genomics Europe (BGE) project, supporting ERGA’s aims of promoting transnational cooperation to promote advances in the application of genomics technologies to protect and restore biodiversity (Mazzoni et al. 2023).

## Materials & Methods

ERGA’s sequencing strategy includes Oxford Nanopore Technology (ONT) and/or Pacific Biosciences (PacBio) for long-read sequencing, along with Hi-C sequencing for chromosomal architecture, Illumina Paired-End (PE) for polishing (i.e. recommended for ONT-only assemblies), and RNA sequencing for transcriptomic profiling, to facilitate genome assembly and annotation.

### Sample and Sampling Information

A female specimen of *Graellsia isabellae* was collected by Marta Vila and Neus Marí-Mena at the type locality (Peguerinos, Ávila, Castilla y León, Spain) on 25 June 2022. In addition, two male (homogametic) individuals of the same species were sampled. The first one was also collected on 25 June 2022 by Vila & Marí-Mena at Peguerinos. The second one was collected by Roger Vila in Casa de la Collada (Collsuspina, Catalunya, Spain) on 09 June 2023. The sampling of all specimens was performed under permission AUES/CYL/168/2022 issued by the Junta de Castilla y León, and SF/0187/23 issued by the Generalitat de Catalunya, Spain. The female was attracted to a light trap, and male attraction was additionally strengthened by the use of synthetic female pheromones (Millar et al. 2010). Specimens were subsequently collected using a butterfly net and identified based on external morphology. Specimens were preserved in dry ice until arrival at the lab where the specimens were later preserved at −80°C in a freezer.

### Vouchering information

Physical reference materials for the male sampled by Roger Vila have been deposited in the Museo Nacional de Ciencias Naturales (Entomology Collection) Madrid, Spain https://www.mncn.csic.es under the accession number MNCN:Ent:372276.

Frozen reference somatic tissue material is available from a proxy voucher at the Biobank of the Museo Nacional de Ciencias Naturales https://www.mncn.csic.es under the voucher ID MNCN:ADN:151.721. This specimen was non-lethally sampled in 2008 at the type locality by Marta Vila and Carlos Lopez-Vaamonde under permission of Junta de Castilla y León (EP/CYL/225/2008).

### Data Availability

*Graellsia isabellae* and the related genomic study were assigned to Tree of Life ID (ToLID) ‘ilGraIsab1’ and all sample, sequence, and assembly information are available under the umbrella BioProject PRJEB77895. The sample information is available at the following BioSample accessions: SAMEA114400479 and SAMEA116288378. The genome assembly is accessible from ENA under accession number GCA_964265105.1 and the annotated genome is available through the Ensembl Beta page (https://beta.ensembl.org/). Sequencing data produced as part of this project are available from ENA at the following accessions: ERX12756251, ERX13166513, ERX13166514, and ERX13166515. Documentation related to the genome assembly and curation can be found in the ERGA Assembly Report (EAR) document available at https://github.com/ERGA-consortium/EARs/blob/main/Assembly_Reports/Graellsia_isabellae/ilGraIsab1/ilGraIsab1.1_EAR.pdf. Further details and data about the project are hosted on the ERGA portal at https://www.ebi.ac.uk/biodiversity/organism/SAMEA114400476.

### Genetic Information

The estimated genome size, based on ancestral taxa, is 0.65 Gb. This is a diploid genome with a haploid number of 31 chromosomes (2n = 62). Males are expected to be homogametic with regard to sex chromosomes (ZZ), while females are yet to be confirmed as ZW or Z0 (ancestral W chromosome has been reported to fuse with an autosome in some *Saturniidae* species (Yoshido et al. 2011). All information for this species was retrieved from Genomes on a Tree (Challis et al. 2023).

### DNA/RNA processing

DNA was extracted from the head and thorax using the Blood & Cell Culture DNA Mini Kit (Qiagen), following the manufacturer’s instructions. DNA quantification was performed using a Qubit dsDNA BR Assay Kit (Thermo Fisher Scientific) and DNA integrity was assessed using a Genomic DNA 165 Kb Kit (Agilent) on the Femto Pulse system (Agilent). The DNA was stored at +4°C until used.

RNA was extracted using an RNeasy Mini Kit (Qiagen) according to the manufacturer’s instructions. RNA was extracted from three different specimen parts: legs, head, and abdomen. RNA quantification was performed using the Qubit RNA BR kit and RNA integrity was assessed using a Fragment Analyzer system (RNA 15nt Kit, Agilent). The RNA was pooled in a 1:2:2 ratio (leg:head:abdomen) for the library preparation and stored at −80°C until used.

### Library Preparation and Sequencing

For long-read whole genome sequencing, a library was prepared using the SQK-LSK114 Kit (Oxford Nanopore Technologies, ONT), which was then sequenced on a PromethION 24 A Series instrument (ONT). A short-read whole genome sequencing library was prepared using the KAPA Hyper Prep Kit (Roche). A Hi-C library was prepared from head and thorax using the ARIMA High Coverage Hi-C Kit (ARIMA), followed by the KAPA Hyper Prep Kit for Illumina sequencing (Roche). The RNA library was prepared from the pooled sample using the KAPA mRNA Hyper prep kit (Roche). All the short-read libraries were sequenced on a NovaSeq 6000 instrument (Illumina).

In total 265x Oxford Nanopore, 78x Illumina WGS shotgun, and 63x HiC data were sequenced to generate the assembly.

### Genome Assembly Methods

The genome was assembled using the CNAG CLAWS pipeline (Gomez-Garrido 2024). Briefly, reads were preprocessed for quality and length using Trim Galore v0.6.7 and Filtlong v0.2.1, and initial contigs were assembled using NextDenovo v2.5.0, followed by polishing of the assembled contigs using HyPo v1.0.3, removal of retained haplotigs using purge-dups v1.2.6 and scaffolding with YaHS v1.2a. Finally, assembled scaffolds were curated via manual inspection using Pretext v0.2.5 with the Rapid Curation Toolkit (https://gitlab.com/wtsi-grit/rapid-curation) to remove any false joins and incorporate any sequences not automatically scaffolded into their respective locations in the chromosomal pseudomolecules (or super-scaffolds). Finally, the mitochondrial genome was assembled as a single circular contig of 15,247 bp using the FOAM (https://github.com/cnag-aat/FOAM) pipeline and included in the released assembly (GCA_964265105.1). Summary analysis of the released assembly was performed using the ERGA-BGE Genome Report ASM Galaxy workflow (https://workflowhub.eu/workflows/1103?version=3).

## Results

### Genome Assembly

The genome assembly has a total length of 560,920,942 bp in 32 scaffolds (Z chromosome included) and the mitogenome (Figures 1 & 2), with a GC content of 35.6%. The assembly has a contig N50 of 18,864,205 bp and L50 of 14 and a scaffold N50 of 20,417,927 bp and L50 of 13. The assembly has a total of 6 gaps, totaling 1.2 kb in cumulative size. The single-copy gene content analysis using the Lepidoptera database with BUSCO v5.5.0 (Manni et al. 2021) resulted in 98.2% completeness (97.9% single and 0.3% duplicated). 91.9% of reads k-mers were present in the assembly and the assembly has a base accuracy Quality Value (QV) of 48.5 as calculated by Merqury (Rhie et al. 2020).

**Figure 1.**
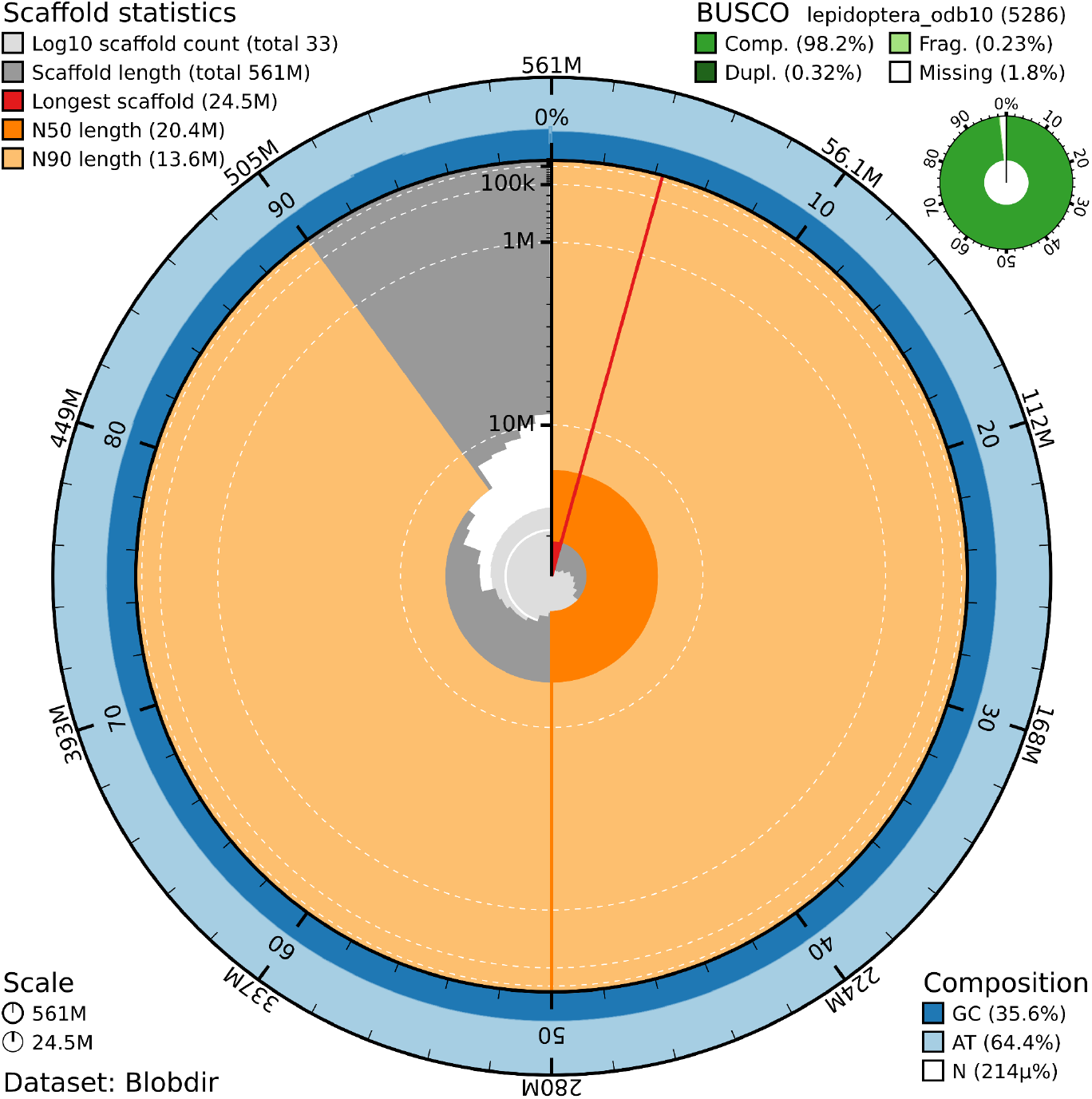
Snail plot summary of assembly statistics. The main plot is divided into 1,000 size-ordered bins around the circumference, with each bin representing 0.1% of the 560,920,942 bp assembly including the mitochondrial genome. The distribution of sequence lengths is shown in dark grey, with the plot radius scaled to the longest sequence present in the assembly (24.5 Mb, shown in red). Orange and pale-orange arcs show the scaffold N50 and N90 sequence lengths (20,417,927 bp and 9,853,470 bp), respectively. The pale grey spiral shows the cumulative sequence count on a log-scale, with white scale lines showing successive orders of magnitude. The blue and pale-blue area around the outside of the plot shows the distribution of GC, AT, and N percentages in the same bins as the inner plot. A summary of complete, fragmented, duplicated, and missing BUSCO genes found in the assembled genome from the Lepidoptera database (odb10) is shown in the top right.

**Figure 2.**
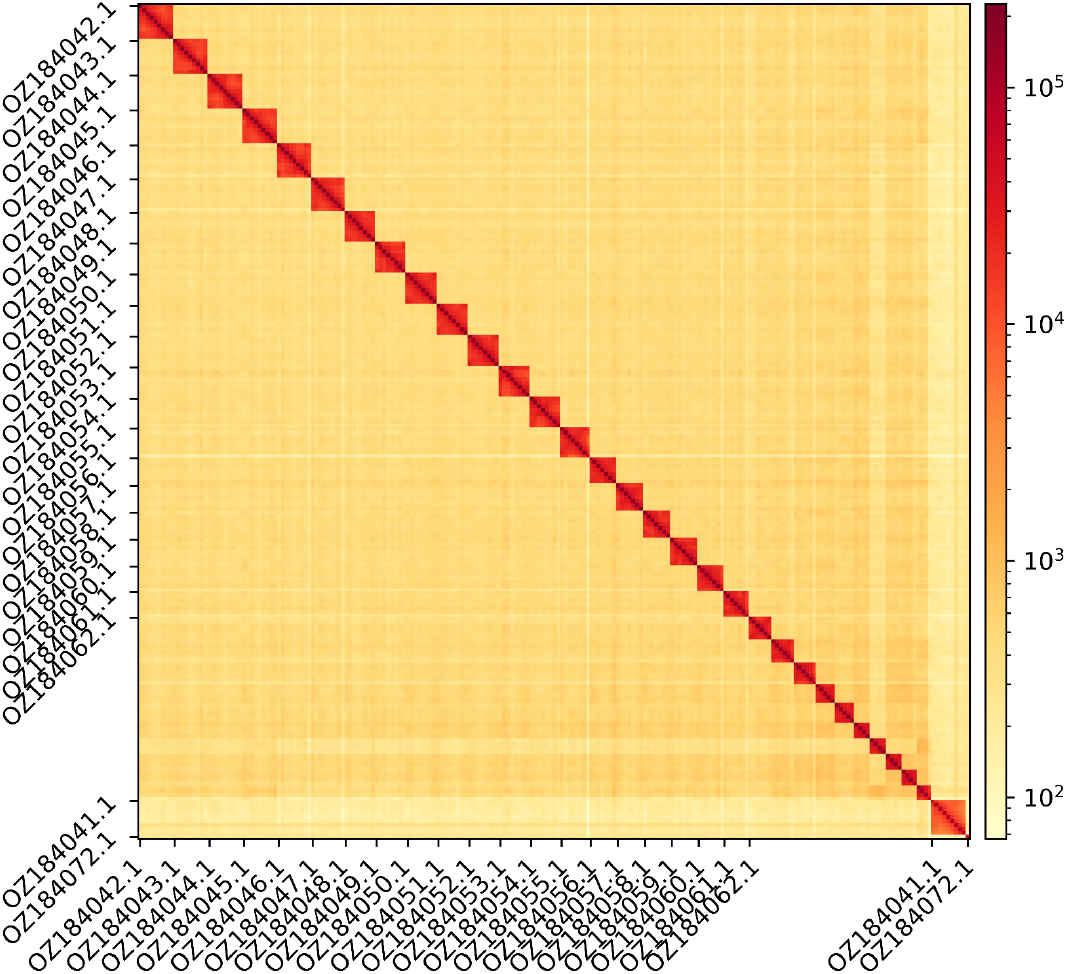
Hi-C contact map showing spatial interactions between regions of the genome. The diagonal in the Hi-C contact map generated with HiCExplorer corresponds to the intra-chromosomal contacts, depicting chromosome boundaries. The frequency of contacts is shown on a logarithmic heatmap scale. Hi-C matrix bins were merged into a 25 kb bin size for plotting. Due to space constraints on the axes, only the GenBank names of the 21st largest autosomes, the Z chromosome (GenBank name: OZ184041.1), and the mitochondrial genome (GenBank name: OZ184072.1) are shown.

## Acknowledgements

We would like to express our gratitude to José Sevilla, Arnau Sevilla, and Telmo Romero, for their invaluable assistance during fieldwork. We acknowledge the assembly reviewer, Sarah Pelan, from the Wellcome Sanger Institute.

## Author Contributions

MV coordinated the project; MV, NM-N, and RV collected the species; MV, NM-N, and RV identified the species; DG-S, CL-V, and AR obtained the barcode sequences; MV and RV sampled and preserved biological material and provided metadata; RM, AsB, RO, THS, and TM provided support in sampling, shipping of biological material, metadata collection, and management; LA and MG extracted DNA, prepared libraries, and performed sequencing; FCF, JGG and FC performed genome assembly and curation under the supervision of TA; CB generated the analysis and report. All authors contributed to the writing, review, and editing of this genome note and read and approved the final version.

